# Cystic Fibrosis Acidic Microenvironment Determines Antibiotic Susceptibility and Biofilm Formation of *Pseudomonas aeruginosa*

**DOI:** 10.1101/2020.10.15.339978

**Authors:** Qiao Lin, Joseph M. Pilewski, Y. Peter Di

**Affiliations:** Department of Environmental and Occupational Health, University of Pittsburgh, Pittsburgh, Pennsylvania, USA; Pulmonary, Allergy, and Critical Care Medicine Division, University of Pittsburgh Medical Center, Pittsburgh, Pennsylvania, USA

**Keywords:** cystic fibrosis, Pseudomonas aeruginosa, antibiotic resistance, bacterial evolution, acidic pH

## Abstract

*Pseudomonas aeruginosa* is the most prevalent bacterial species that contributes to cystic fibrosis (CF) respiratory failure. The impaired function of cystic fibrosis transmembrane conductance regulator leads to abnormal epithelial Cl^−^ / HCO_3_^−^ transport and acidification of airway surface liquid. However, it remains unclear why *Pseudomonas aeruginosa* preferentially colonizes in the CF lungs. In this study, we carried out studies to investigate if lower pH helps *Pseudomonas aeruginosa* adapt and thrive in the CF-like acidic lung environment. Our results reveal that *Pseudomonas aeruginosa* generally forms more biofilm and induces antibiotic resistance faster in acidic conditions and that this can be reversed by returning the acidic environment to physiologically neutral conditions. *Pseudomonas aeruginosa* appears to be highly adaptive to the CF-like acidic pH environment. By studying the effects of an acidic environment on bacterial response, we may provide a new therapeutic option in preventing chronic *Pseudomonas aeruginosa* infection and colonization.

## Introduction

Cystic fibrosis (CF) is a genetic disease involving compromised function of cystic fibrosis transmembrane conductance regulator (CFTR) that leads to impaired airway host defense and therefore causes lung inflammation and bacterial infections (1, 2). The environment in CF lungs are thought to be acidic as evident by the lower-than-neutral pH value of the airway surface liquid (ASL) in newborn CF pigs, as well as in differentiated human and porcine primary epithelial cell cultures, compared to non-CF controls (1, 2). The malfunction of CFTR also leads to elevated Na^+^ and Cl^−^ levels in the airway, which inhibits the natural antimicrobial factors in ASL (3). *Pseudomonas aeruginosa* (*P. aeruginosa*) is an opportunistic gram-negative pathogen that is the most dominant bacterial species in adult CF and contributes to the associated high mortality rates (4). The prevalence of *P. aeruginosa* lung infection gradually increases over time from approximately 20% to 70% in CF patients from age 2 to 45 (5), which coincides with an increase in CF disease severity. It is noted that the bacterial biofilm formation in the CF airway likely contributes to the chronic colonization of *P. aeruginosa* (6). Additionally, it has been reported that extracellular DNA (eDNA) is the most abundant polymer within the *P. aeruginosa* biofilm matrix (7, 8). Previous studies suggested that eDNA acidifies *P. aeruginosa* biofilm and promotes resistance to aminoglycoside antibiotics (9). The CF lung is also burdened with acidic eDNA originated from the immune cells recruited by the chronic bacterial infections (10). In this study, we sought to determine the effects of CF-like acidic environment on the pathogenic ability of *P. aeruginosa*. Our results indicated that the acidic environment stimulates increased *P. aeruginosa* biofilm formation, promotes faster adaption of bacteria towards elevated antibiotic resistance, and increases expressions of multiple biofilm/virulence-related genes. Furthermore, the acidic environment on the apical surface of differentiated primary bronchial epithelial cells isolated from CF patients (CFBEs) results in increased *P. aeruginosa* attachment and bacterial numbers than the physiologically neutral environment on the surface of cultured primary human bronchial epithelial cells from normal subjects (HBEs). These adverse effects of a CF-like acidic environment can be ameliorated by modulating the acidic environment into physiologically neutral conditions. The varying characteristics and behavior of *P. aeruginosa* in different pH conditions may provide additional treatment targets and options for CF sufferers in preventing chronic *P. aeruginosa* infection and colonization.

## Results

### Response of *P. aeruginosa* to CF-like microenvironment

Bacteria expand their population in the natural environment and in the host via two different forms. The planktonic form of free-moving bacteria is the typical way bacteria spread themselves at the initial stage of reaching a new environment. In contrast, the accumulated form of bacteria, so-called “biofilm”, represents another important pathogenic mechanism. *P. aeruginosa* gradually increase their presence in the CF airways after the pathological phenotypes of CFTR malfunction appear more obviously with increased age of the CF sufferers (5). It is well-documented that compromised functions of CFTR and the CFTR-modulated HCO_3_^−^ secretion in CF result in abnormal environment of higher salt (~100±5mM) (3) and lower pH (~6.7±0.3) (1, 2, 11) in the CF ASL than those of normal subjects.

We first examined the effects of salt concentration and acidic condition on bacterial proliferation in both planktonic and biofilm mode of growth in order to understand the effects of CF-like microenvironment on the behavior of *P. aeruginosa*. Two widely used *P. aeruginosa* lab strains (PAO1 and PA14) and two multi-drug resistant (MDR) *P. aeruginosa* clinical strains isolated from CF patients (*P.a.* 129-5 and *P.a.* 152-19) were selected for this experiment. The growth curves indicated that various salt concentrations (addition of 50, 100 and 150mM NaCl) and pH conditions (pH= 6.0, 6.5, 7.0 and 7.5 adjusted by HCl) did not result in any noticeable difference on the proliferation rate of planktonic *P. aeruginosa* in all four strains (Fig. S1). There was less biofilm biomass formed with the increasing NaCl concentrations in PAO1 while the effect of salt concentrations on biofilm formation was not apparent in the other three *P. aeruginosa* strains (Fig. S2). Nonetheless, we observed significant changes on the initial attachment of bacterial biofilm under acidic pH conditions after just 3 hours of incubation (Fig. S2). Except PA14, all the other three tested *P. aeruginosa* strains showed similar results of increased biofilm formation at acidic (pH<7) conditions. These results suggest that the acidic environment likely promoted the initial attachment of *P. aeruginosa*.

### Increased *P. aeruginosa* antibiotic resistance under acidic pH conditions

It is known that ineffective treatments of antibiotics such as inhaled tobraomycin and ceftazidime in CF respiratory infections, subsequently lead to lifelong *P. aeruginosa* infection (5, 12). Thus, we sought to determine if the CF-like acidic environment enhanced bacterial resistance to antibiotics. Three different classes of commonly used standard-of-care antibiotics, including ceftazidime (β-lactam), ciprofloxacin (fluoroquinolone) and tobramycin (aminoglycosides) each with a distinct antibacterial mechanism, were tested against the same four bacterial strains of *P. aeruginosa* at different pH conditions. Our results indicated that acidic pH environment alone does not notably change the *P. aeruginosa* proliferation rate (Fig. 1A). However, acidic conditions increased the minimum inhibitory concentration (MIC) of all three clinically used antibiotics as demonstrated in the elevated growth curves compared to pH 7.5 when the same dosage of each antibiotic was used in all varying pH conditions (Fig. 1B). In all cases, *P. aeruginosa* grown under the acidic conditions (pH=6.0 & pH=6.5) demonstrated the elevated resistance to the antibiotic treatments as their growth curves were closer to those of the “no antibiotic treatment” controls. The slope of the growth curve gradually decreases as the pH value increases. To determine if the increased bacterial growth in lower pH conditions was due to permanent degradation or temporary inactivation of antibiotics in the acidic environment, we pre-incubated the antibiotics in physiologically neutral pH 7.5 or acidic pH 6.0 for 5 hours prior to the bacterial growth kinetic experiments. Interestingly, the antibiotics pre-exposed to pH 6.0 and pH 7.5 showed similar growth inhibition curves when the growth inhibition assay (GIA) was performed at the neutral pH 7.5 condition (Fig. S3). The results indicated that antibiotics pre-exposed to acidic conditions (pH=6.0) did not lose their drug potency and regained their antibacterial activity after returning to the physiologically neutral condition (pH=7.5). These data suggested that *P. aeruginosa* could be more resistant to antibiotic treatment in an acidic environment.

**Fig. 1.**
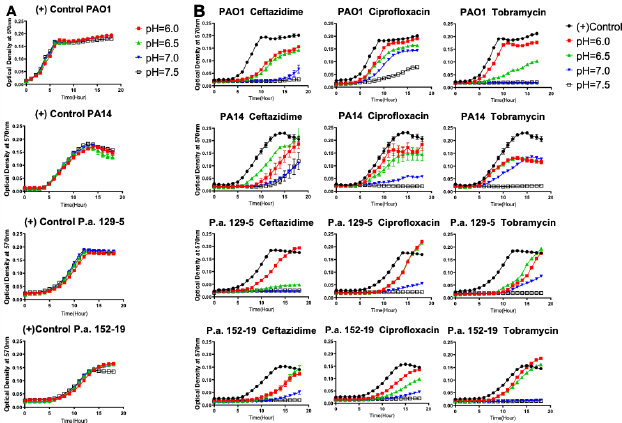
Acidic pH conditions impair the antimicrobial activity of standard-of-care antibiotics against *P. aeruginosa*. (**A**) *P. aeruginosa* growth curve in pH adjusted, antibiotic-free medium; **(B)** *P. aeruginosa* growth curve in pH adjusted medium, supplemented with ceftazidime, ciprofloxacin and tobramycin at the respective MIC for each antibiotic. Optical density at 570nm was measured every hour for 18 hours at 37°C in a microplate reader. Results are mean ± SEM from three independent experiments with bacteria grew in duplicates for each condition.

To further explore if acidic pH promotes *P. aeruginosa* antibiotic resistance, we carried out studies to investigate bacterial adaption (evolution) to antibiotics in the acidic and neutral pH conditions. Both planktonic and biofilm mode of *P. aeruginosa* were grown under the treatment of ceftazidime, ciprofloxacin and tobramycin, three of the most commonly used antibiotics in treating CF respiratory infection. The *P. aeruginosa* biofilm was generated by growing PA14 on acrylic beads and transferred daily to a new culture tube with a new sterile bead in the existence of antibiotics (Fig. 2A). The planktonic culture served as a control to the biofilm evolution model in contrasting the different growth modes (Fig. 2B). The new MIC of each PA14 culture condition was determined by the survival of each population after 24 hours of incubation in a fresh medium containing doubled antibiotic concentration on a daily basis. After 15 days of continuous growth and evolution, PA14 populations acquired increased levels of resistance against all three antibiotics. The acidic environment significantly increased the MIC required for the tested antibiotics to kill *P. aeruginosa* (Fig. 2C-E), regardless of the antibiotic killing mechanisms. These results strongly indicate that subtle pH change under pathophysiological conditions determines the magnitude and timing *P. aeruginosa* adapts to antibiotic selection pressure. The reference strain of *P. aeruginosa* (PA14) was selected because we sought to focus on investigating the pH effect without additional concerns of horizontally transferred plasmids or bacteriophages which could exist in clinical strains. Additionally, biofilm formation of PA14 is not significantly influenced by pH variations (Fig. 3, Fig. S2). Therefore, the changes of MIC in this bacterial evolution model is less likely influenced by pH modulated biofilm formation. Interestingly, in contrast to the PA14 ancestor, evolved PA14 progeny bacteria demonstrated indisputable ability to survive and adapt to highly stressful antibiotic treatments, while acidic pH significantly expedited this process.

**Fig. 2.**
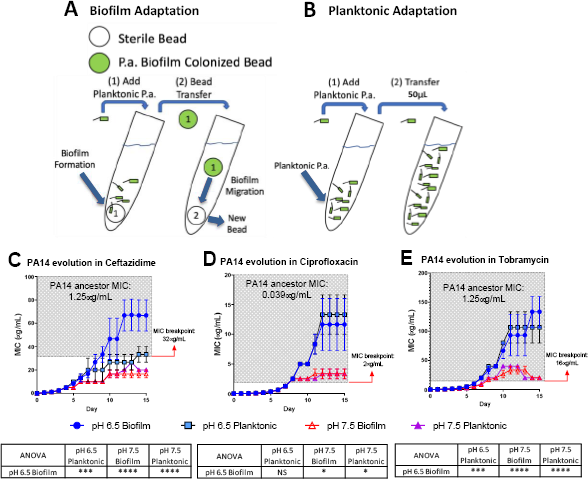
Acidic pH promotes faster accumulation of adaptive resistance of *P. aeruginosa* against antibiotics compared to pH 7.5. (A and B) Schematics of the *P. aeruginosa* PA14 antibiotic adaptation experiment: biofilm on beads (A) and planktonic bacteria (B) were transferred to fresh m63 media (pH=6.5/7.5, adjusted by HCl) every 24 hours; (C)-(E) The PA14 biofilm/planktonic cultures were treated with ½ MIC at day 1. Antibiotic concentrations were doubled after every bead/planktonic transfer (*n*=3). The grey boxes on figures (C)-(E) denote MIC values that are considered antibiotic resistant according to the guideline “MIC Breakpoints for *Pseudomonas aeruginosa*”, published by The Clinical and Laboratory Standards Institute. Data are mean ± SEM. One-way ANOVA was performed by comparing the combined day 14 and 15 MIC values. * *p*<0.05; *** *p*<0.001; **** *p*<0.0001; NS: not significant.

**Fig. 3.**
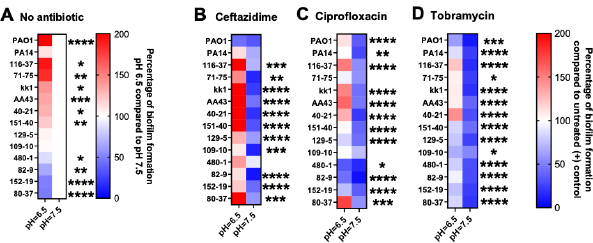
Acidic pH modulates *P. aeruginosa* biofilm formation and impairs antibiotic biofilm prevention against *P. aeruginosa*. All *P. aeruginosa* strains (2 lab strains and 12 clinical strains) were incubated in pH adjusted m63 medium for 18 hours. Crystal violet staining method was used to quantify biofilm formation (*n*=6). (A) *P. aeruginosa* biofilm formation without antibiotic treatment. Biofilm formation in pH 7.5 was served as positive control. (B-D) *P. aeruginosa* biofilm formation treated by ceftazidime, ciprofloxacin and tobramycin at the concentrations of 1x MIC of each *P. aeruginosa* strain (MIC acquired in normal pH 7.5 condition). Percentage of antibiotic treated biofilm formation was calculated by comparing to each untreated *P. aeruginosa* strain in pH 7.5. The color scale bars represent percentage of biofilm formation compared to appropriate control groups. Red scale: increased biofilm formation (100% − 200%); white: no change (100%); blue scale: decreased biofilm formation (0 – 100%). Data were collected from two independent experiments. Unpaired *t* tests were used for statistical analysis between each pH 6.5 and pH 7.5 conditions. * *p*<0.05; ** *p*<0.01; *** *p*<0.001; **** *p*<0.0001; otherwise not significant.

### Acidic conditions promote biofilm formation of *P. aeruginosa* clinical isolates and compromise biofilm prevention activities of antibiotics

To further validate the hypothesis that acidic pH promotes biofilm formation and impairs antibiotic efficacy, we evaluated biofilm formation and antibiotic biofilm prevention under acidic conditions of fourteen *P. aeruginosa* strains. These include twelve clinical *P. aeruginosa* isolates that were originally obtained from CF patients and two standard lab strains (PAO1 and PA14). In Fig. 3A, CF-like, acidic pH 6.5 condition alone stimulated more mature biofilm formation from approximately 71% (10/14) of all *P. aeruginosa* strains compared to pH 7.5. The difference is significant and more noticeable when 1x MIC of ceftazidime, ciprofloxacin and tobramycin were used in *P. aeruginosa* biofilm prevention. In Fig. 3B-D, fourteen *P. aeruginosa* strains were treated using their respective MIC dosage (Table S1) in pH 6.5 and 7.5 conditions. The majority of antibiotic treated *P. aeruginosa* strains formed more biofilm in the pH 6.5 condition compared to the pH 7.5 condition. These biofilm data are in accordance with our growth kinetic studies and the PA14 evolution studies (Fig. 1 and 2). All experiments demonstrated a similar pattern: that ceftazidime, ciprofloxacin and tobramycin are generally less effective in preventing *P. aeruginosa* growth and biofilm formation at the acidic pH conditions.

### Neutralization of acidic pH in CF epithelial cells restores impaired host defense

Acidic pH impairs important host defense mechanisms such as the ASL antibacterial activities of antimicrobial peptides including β-defensin-1, −3 and LL-37 (13–15). Our results demonstrate that the acidic pH in the CF-like microenvironment increases bacterial resistance against antibiotics and enhances biofilm formation, which may promote *P. aeruginosa* colonization. We further determined if the neutralization of the acidic ASL of differentiated human bronchial epithelial cells that were maintained under air-liquid interface (ALI) culture could help alleviate *P. aeruginosa* infection (Fig. 4A). The primary human bronchial epithelial (HBE, non-CF) cells and CF bronchial epithelial (CFBE) cells were used. We tested ouabain, an ATP12A inhibitor that inhibits H^+^ secretion by human epithelial cells, thus raising the pH value of cultured epithelial cells (2). Without any treatment, the ASL of cultured CFBE cells was more acidic than non-CF HBE cells (Fig. 4B). However, ouabain successfully elevated pH value of CFBE ASL from approximately 6.7 to 7.6. The increased pH value reflected neutralized ASL of CFBE, which consequently resulted in decreased CFU of *P. aeruginosa* (PAO1) grew on top of human epithelial cells (Fig. 4C). The ouabain modulation of pH in ASL of HBE was minimal, which did not result in significant changes of *P. aeruginosa* CFU (Fig. 4B). Ouabain itself does not display any bactericidal activity at the treatment concentration of 20μM (Fig. S4).

**Fig. 4.**
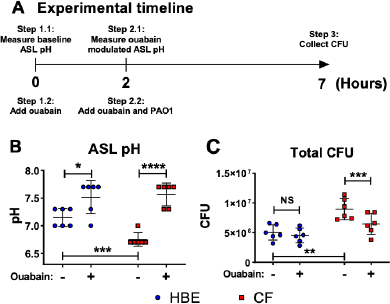
Ouabain helps restoring host defense activities by reversing the acidic pH to neutral pH in differentiated human bronchial epithelial cell cultures. (**A**) Experimental timeline. pH values were measured on the apical side of air-liquid interface cultured cells before and after two hours of treatment with/without 20μM ouabain; PAO1 was then added to the apical side of cultured cells with ouabain or DMSO solvent control for additional 5 hours. (**B**) Effect of ouabain treatment on pH values in non-CF (HBE) and CF epithelial cells. The change of pH in epithelial apical wash was measured by narrow range pH test strips (pH 6-7.7, resolution 0.3 pH unit; n=6). (**C**) Effect of ouabain treatment on *P. aeruginosa* CFU in non-CF (HBE) and CF epithelial cells. PAO1 was incubated on the apical side of cultured HBE and CF epithelial cells in the existence of 20μM ouabain or DMSO (solvent control). Biofilm and planktonic CFU were obtained by plated on agar plates for total PAO1 CFU (n=6). Data were collected from two independent experiments. Unpaired *t* tests were used for statistical analysis. Data are mean ± SEM. * *p*<0.05; ** *p*<0.01; *** *p*<0.001; **** *p*<0.0001; NS: not significant.

### Acidic conditions increase the expression of biofilm-related genes

To evaluate if acidic microenvironment-enhanced *P. aeruginosa* infection in CF is regulated through increased bacterial biofilm formation, we determined the gene expression of a panel of biofilm/virulence-related genes (Fig. 5) in association with the observed biomass changes. Multiple biofilm-related genes such as *tolA, ndvB, rhlA*, and *rhlB* were all significantly increased in the acidic conditions of biofilm formation.

**Fig. 5.**
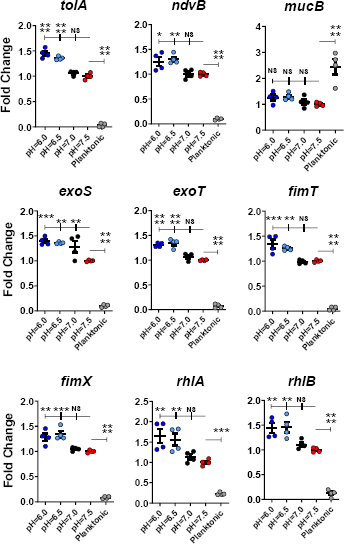
Acidic pH conditions increase biofilm/virulence-related gene expression of *P. aeruginosa*. *P. aeruginosa* PAO1 biofilm was cultured in different pH conditions for 24 hours. Physiologically related condition in pH 7.5 was used as a control for biofilm growth. Planktonic form of *P. aeruginosa* inoculated in pH 7.5 was also included for comparison (n=4). Data were representative of three independent experiments. Results are mean ± SEM. One-way ANOVA and Dunnett’s multiple comparisons test were used for statistical analysis. * *p*<0.05; ** *p*<0.01; *** *p*<0.001; **** *p*<0.0001; NS: not significant.

Additionally, the expression of several virulence-associated genes such as *exoS, exoT, fimT*, and *fimX* were also significantly increased in acidic environments (pH 6.0 and 6.5) compared to in physiological conditions (pH 7.0 and 7.5). Interestingly, the expression of the planktonic-associated *mucB* gene did not vary among all acidic and physiological biofilm-forming pH conditions. However, it differed significantly between the biofilm and planktonic forms of *P. aeruginosa* (Fig. 5).

## Discussion

*P. aeruginosa* is the most dominant bacterial pathogen associated with CF disease severity and mortality. The CF microenvironment resulted from malfunction of CFTR may have unwanted effect in contributing to the infection and colonization of *P. aeruginosa*. However, the underlying pathogenic mechanisms remain to be elucidated. In this study, we investigated the CF-like acidic microenvironment on the behavior of *P. aeruginosa* and demonstrated that multiple *P. aeruginosa* clinical isolates significantly increased their biofilm formation and antibiotic resistance under acidic conditions. Importantly, this pathological factor of acidic pH also activated a series of biofilm- and virulence-related genes of *P. aeruginosa*. Under the selection pressure in an experimental condition with gradually increased antibiotic concentrations, we demonstrated that *P. aeruginosa* adapted to the antibiotics quickly and evolved much faster in pH 6.5 than in pH 7.5, which eventually acquired strong antibiotic resistance in only 15 days. These finding suggest that the acidic CF microenvironment likely play a critical role in facilitating the colonization of *P. aeruginosa* despite the antibiotic treatment.

It has been suggested that the lowered airway pH is associated with impaired host defense mechanisms (1, 3, 13–17). By neutralizing the acidic environment through inhibiting ATP12A, the pH value increased on the apical side of the differentiated human primary airway epithelial cells (CFBEs and non-CF HBEs). The increased pH not only could potentially restore the antimicrobial activity of the naturally secreted host defense factors by airway epithelial cells such as SPLUNC1 (18, 19), but also may prevent biofilm formation and/or pH-induced drug-resistance of *P. aeruginosa*. Interestingly, *P. aeruginosa* appears to be the only bacterial species that displayed significantly increased biofilm formation under CF-like acidic environment among the notorious MDR ESKAPE pathogens (Fig. S5), which include *Enterococcus faecium, Staphylococcus aureus, Klebsiella pneumoniae, Acinetobacter baumannii, Pseudomonas aeruginosa*, and *Enterobacter spp.* Unlike the other five ESKAPE bacterial strains, *P. aeruginosa* alkalinizes the environment when cultured in minimal medium (m63 supplemented with L-arginine). After overnight inoculation, *P. aeruginosa* medium pH generally increases from 7.5 to 8.0 (data not shown), which suggests increased proton uptake or ammonia production. Our data provide an important link between the worsened CFTR function-associated acidic microenvironment and enhanced *P. aeruginosa* biofilm formation supported by the evidence of *P. aeruginosa* biofilm-related phenotypic/genetic changes and multiple microenvironment acidification factors.

The unique responses of *P. aeruginosa* to acidic environments provide potential therapeutic insights. First, *P. aeruginosa* can take advantage of the acidic CF-like lung microenvironment and significantly increased their biofilm biomass especially under common antibiotic treatments (Fig. 3). Second, there are a number of means to neutralize the acidic CF lung environment. By inhibiting ATP12A, the secretion of H^+^ is inhibited and therefore pH is elevated (Fig. 4B). In addition, the extracellular H^+^ could also be neutralized by commonly used substances, such as sodium bicarbonate (9, 20) or hydroxide salts. The increased bacterial susceptibility to host factors after raising the acidic pH to neutral pH suggests an alternative approach in treating CF chronic infection induced by *P. aeruginosa*. Recent studies showed evidence that CFTR modulator ivacaftor increases the function of several genotypes of human CFTR mutations. The increased CFTR activity reduces sweat Cl^−^, which may subsequently result in elevated ASL pH (21).

In addition to the efforts to neutralize the acidic host environment, it is also important to look into the complex response of *P. aeruginosa* to acidic pH and how this response leads to antibiotic resistance. Our gene expression studies provided a mechanistic insight of pH-related pathways in pathogenic *P. aeruginosa*. Acidic conditions at pH values of 6 and 6.5 significantly increased the expression of *tolA*, which is activated in biofilms (22) and its product is responsible for aminoglycoside-resistance in *P. aeruginosa* by decreasing its permeability and blocking the entrance of antibiotics (23–27). Cyclic glucans are circular polymers of glucose that have been shown to help *P. aeruginosa* adapt to low osmotic media and flagella-mediated twitching mobility (28). Cyclic glucans also have the ability to physically interact with tobramycin and therefore eliminate antibiotics before they could reach their targeted site of action (29). The *ndvB* gene encodes for a glucosyltransferase that is required for the synthesis of cyclic-b-(1, 3)-glucans (30). At pH 6.0 and 6.5 conditions, there was a noticeably higher *ndvB* gene expression in *P. aeruginosa* biofilm compared to the pH 7.5 of physiological control. Planktonic form of PAO1 expressed significantly less *ndvB* than any biofilm group. The elevated expression of *ndvB* could be one of the factors that contributes to the acidic pH-induced antibiotic resistance in *P. aeruginosa* (Fig. 1-4).

Both *rhlA* and *rhlB* are required for rhamnolipids synthesis (31), a biosurfactant that contributes to the swarming motility of *P. aeruginosa* in order to promote biofilm colonization. Rhamnolipids are known to interfere with phagocytosis (32) and normal tracheal ciliary function (33). The expression of these virulence factors was also increased under the acidic pH environment. Both *fimT* and *fimX* are required for the biogenesis and functioning of type IV pili and twitching mobility in *P. aeruginosa* (34). These virulence factors are required for biofilm formation (35, 36). Elevated expression of both *fimT* and *fimX* likely associates with stronger ability in biofilm formation and host colonization enhanced under acidic pH conditions. *P. aeruginosa* injects cytotoxins (e.g. ExoS and ExoT) into host cells by utilizing the type III secretion system (37). Acidic pH conditions activated *exoS* and *exoT* will likely lead to delayed wound healing, impaired phagocytosis and spread of *P. aeruginosa* (38).

The *mucB* gene is a negative regulator of the sigma factor AlgU. Inactivation of this gene will convert *P. aeruginosa* into alginate-producing mucoid forms (39, 40). All biofilm groups have reduced *mucB* expression compared to the planktonic *P. aeruginosa*. This suggests that the free-swimming planktonic *P. aeruginosa* are less mucoid than their biofilm counterparts. Unlike all other biofilm-related genes, acidic pH did not stimulate a significant change of *mucB* expression in *P. aeruginosa* biofilms. At the early stages of bacterial colonization, alginate is usually not considered to be a major component of the *P. aeruginosa* biofilm extracellular matrix (41).

Previous studies demonstrated that when *P. aeruginosa* is under acidic stress, the bacterial outer membrane permeability could decrease and result in decreased antibiotic uptake. The decrease of bacterial outer membrane permeability (PhoPQ/PmrAB-controlled surface modifications) is pH-mediated (9, 17). This partly explains why acidic pH induces instant but moderate antibiotic resistance, approximately 2 to 8-fold increase (Supplementary Table 1). In addition, acidic pH likely modulates aminoarabinose-modified LPS and spermidine, which mask bacterial negative surface charges and block the entrance of aminoglycosides (9, 42, 43). Our results demonstrated that *P. aeruginosa* grew under other classes of antibiotics (β-lactam and fluoroquinolone), in addition to the previously published aminoglycosides (9), also increase the acquired antibiotic resistance. These outcomes may subsequently contribute to the chronic colonization of *P. aeruginosa* in CF, which is the major cause of the worsening lung function in CF. While the previous publications focused on selected genetic mutations associated with antibiotic resistance, our studies aimed at examining the CF-like acidic microenvironment in regulating *P. aeruginosa* biofilm formation, and how does acidic pH promote faster bacterial evolution in developing resistance against antibiotics of three different killing mechanisms (Fig. 2, 50 to 300-fold MIC increase in acidic populations after 15 days of continuous evolution).

There is no consensus on whether or not the CF ASL is substantially more acidic than those in normal subjects. We have been able to consistently obtain more acidic pH measurements in human ALI bronchial epithelial cell culture derived from ex-planted lungs of CF patients than those from non-CF subjects (Fig. 4B). However, some studies reported that they were unable to measure significant pH differences in ASL of children w/wo CF (44) and the pH values are highly dependent on the pH probe materials and locations of the measurement (nasal or tracheal) (45). On the contrary, it is also reported that without the innate proton pump ATP12A activity and airway acidification, CFTR deficient mouse are free from opportunistic respiratory infection (1), which makes airway acidification the deciding factor of murine opportunistic lung infections. Additionally, eDNA originated from bacterial biofilm and human immune cells also contributes to the acidification of CF microenvironments, which is evidenced by acidic pH gradients within *P. aeruginosa* biofilms (46–48) and acidified CF exhaled breath condensate (49, 50), all of which are difficult to be discredited by a direct ASL pH measurement in children, who are usually with less prominent CF phenotype and severity. It is likely that the CF microenvironment is much more complicated than what can be currently measured on human subjects. Our results are representative of the effects of acidic environment on *P. aeruginosa* but do not address all abnormal CF conditions

This study was designed to explore the mechanisms of initial *P. aeruginosa* colonization and how the CF-specific lung microenvironment might contribute to this adverse health outcome. *P. aeruginosa* colonization in respiratory tracts is a chronic process and the bacteria likely adapt and evolve in the CF lungs for decades. Longitudinal studies of the CF *P. aeruginosa* genotypes have illustrated that *P. aeruginosa* undergoes extensive genomic DNA mutations in order to survive in the CF lungs (51). Although multiple mutated genes have been identified, there is no evidence to connect any of the mutations to the important CF-specific, low pH acidic lung microenvironment. A limitation of our study is that we could not identify the specific genetic mutations of *P. aeruginosa* that are resulted from CF-like acidic microenvironment. More studies are needed to target this aim and to explore the role of acidic pH in long-term *P. aeruginosa* evolution/adaption to resolve the problem of the persistent biofilm formation and chronic colonization suffered by CF patients.

## Methods

### Bacterial strains

Clinical *P. aeruginosa* strains were isolated from pediatric CF patients with chronic pulmonary infections at Seattle Children’s Research Institute except the following ones, which were from collection of the CF clinic at Medizinische Hochschule of Hannover, Germany: AA43 and KK1(52). The lab strains used in this study were *P. aeruginosa* PAO1 (ATCC, BAA-47) and UCBPP-PA14 (53).

### Preparation of *P. aeruginosa*

All bacteria were retrieved from −80°C frozen stock and streaked on tryptic soy agar plates. Single colonies were picked and incubated in tryptic soy broth (TSB) overnight at 37°C in an orbital shaker. The overnight culture was diluted 1:5 with fresh TSB and incubated for an additional 2 hours for exponential growth. Bacteria were centrifuged at 2,000g for 5 minutes. The pellet was resuspended in 1ml PBS. To ensure reproducible results, bacterial concentration was adjusted to approximately 10^9^ CFU/mL, optical density (OD500nm) = 0.5±0.01 in a spectrophotometer.

### Bacterial growth inhibition assay (GIA)

All GIA studies were performed in 10% TSB diluted in PBS. The 96-well plate was incubated at 37°C in a microplate reader for 18 hours. Optical density (OD) at 570nm was measured every hour with continuous double orbital shaking at 425 cycles per minute. The pH of the bacterial culture was adjusted to 6.0, 6.5, 7.0 and 7.5 using hydrochloride acid (HCl). The starting bacterial concentration in each treatment group was 10^6^ CFU/mL. All pH-adjusted media were filter-sterilized by a syringe filter unit (pore size: 0.22μm, Millipore SLGP033RS) to ensure sterility.

### Biofilm assay

The crystal violet biofilm staining method developed by O’Toole *et al.* was used in this study with slight modifications (54). All biofilm studies were performed in pH adjusted m63 medium supplemented with 1mM MgSO_4_, 25μM FeCl_3_, 40mM of D-glucose and 4mM of L-glutamine. The combination of D-glucose and L-glutamine allows the m63 to maintain its pH after *P. aeruginosa* overnight incubation. The final bacterial concentrations were 5×10^7^ and 10^6^ CFU/ml for biofilm formation and antibiotic biofilm prevention in different pH, respectively. After 18 hours of biofilm formation in a 37°C incubator (with or without antibiotic treatment), the supernatant was discarded and biofilm attached to the plate was stained with 0.5% crystal violet (20% ethanol + 80% deionized water) solution for 15 minutes. The excessive dye was then rinsed off with water and 95% ethanol was added to release the dye from biofilm. OD values were acquired by a microplate reader at the wavelength of 620nm.

### Bacterial evolution in antibiotics

Evolution studies were carried out using m63 medium (same as biofilm assay). The bead transfer-based biofilm evolution model was described previously (55–57). Briefly, the PA14 ancestor was added to 5mL of pH 6.5/7.5 m63 (media pH adjusted by HCl) with ½ MIC of antibiotic treatment at day 1. After 24 hours, the PA14 biofilm (formed on sterile acrylic bead) was transferred to the next tube of fresh m63 medium with a sterile bead inside. For planktonic culture, 50uL of PA14 overnight culture was transferred to the next tube of fresh m63 medium. The dosage of antibiotics was doubled at the time of each transfer. The groups that survived antibiotic treatment were transferred to the next tube. The groups that could not tolerate the elevated antibiotic concentration were incubated again at the prior concentration without doubling antibiotic concentration.

### Primary airway epithelial cell cultures

Differentiated primary human bronchial epithelial cells were derived from lungs removed at the time of lung transplantation at the Center for Organ Recovery and Education (Pittsburgh, PA, USA). Cells were prepared using previously described methods (58, 59) approved by the University of Pittsburgh Institutional Review Board. Donor primary human CF and non-CF bronchial epithelial cells were first isolated from donor tissues and propagate under submerged cell culture. Upon confluence, epithelial cells were disassociated and seeded onto transwell cell culture plate at approximately 2×10^5^ cells/well (Corning, NY). Epithelial cells were maintained in Bronchial Epithelial Cell Growth Medium (Lonza, Basel, Switzerland). The cell culture was changed to air-liquid interface (ALI) by removing apical medium three days after initial cell seeding and maintained for three weeks for cell differentiation.

### Treatment and host defense activity of primary airway epithelial cell cultures

The 4mM ouabain stock solution was prepared and dissolved in DMSO. Ouabain was diluted to 20μM in normal saline and 20μL of the diluted ouabain or DMSO (vehicle control) was added to the primary cell cultures apically and incubated for 2 hours in 37°C supplied with 5% CO_2_. Primary cultured epithelial cells were washed apically with 100μL of PBS 24 hours before experiment. A total of 10μL of the apical fluid was immediately added on pH test strips. After pH reading, all remaining apical fluid was removed and another 20μL of 20μM ouabain or DMSO was added with 50μL of PAO1 suspended in normal saline (10^7^ CFU/insert). Cells were then incubated for additional 5 hours for biofilm to form. All apical supernatant was collected for determining the CFU of unattached planktonic bacteria. The ALI membrane was subsequently removed from the filter and sonicated in 2mL of PBS for 30 seconds at 80% amplitude by the DPS-20 dual processing system (130?W; PRO Scientific) to disassociate congregated PAO1 biofilm. After sonication, both planktonic and biofilm samples of PAO1 were plated on tryptic soy agar plates for total CFU counting.

### Gene expression

A total of 5 x 10^7^ CFU/mL PAO1 was cultured in pH-adjusted (pH=6.0, 6.5, 7.0 and 7.5) DMEM for three hours in 100×15mm round-petri dishes at 37°C. For biofilm RNA samples, supernatant in all petri dishes was discarded and a thin layer of PAO1 biofilm was scraped off using a sterile cell scraper. The same amount of PAO1 was incubated separately in 5 mL of pH 7.5 DMEM at the same time and served as a planktonic control. Total biofilm and planktonic RNA was extracted as previously published (60–62). The cDNA was synthesized by High-Capacity cDNA Reverse Transcription Kit (Applied Biosystems). Gene expression results were obtained using Fast SYBR Green Master Mix (Applied Biosystems) and 7900HT Fast Real-Time PCR System (Applied Biosystems). ΔΔ*C*_t_ values were calculated and analyzed as method previously published (63). The list of primers can be found in Table S2.

### Statistical analysis

Error bars represent mean ± standard error of the mean (SEM). One-way Analysis of Variance (ANOVA) and Tukey’s multiple comparisons test were used to assess the overall change of MIC in the bacterial evolution model. One-way ANOVA and Dunnett’s multiple comparisons test were used to assess the change of bacterial gene expressions among various pH conditions. Two-way ANOVA and Dunnett’s multiple comparisons test were used to assess the effects of pH and salt on bacterial biofilm formation. Student’s t tests were used to assess statistical significance between two subjects, such as biofilm formation and antibiotic biofilm prevention in different pH, human ALI ASL pH and CFU with or without ouabain treatment. * *p*<0.05; ** *p*<0.01; *** *p*<0.001; **** *p*<0.0001; otherwise not significant (NS).

## Data availability

All data generated or analyzed during this study are included in this published article (and its supplementary information files).

## Acknowledgments

This work was supported by the National Institutes of Health (R01 AI-133351 to Y.P.D.). The funding agencies had no role in study design, data collection and analysis, decision to publish, or preparation of the manuscript. We thank Dr. Michael M. Myerburg for providing the primary human CF and non-CF bronchial epithelial cells.

## Author Contributions

Conceptualization, Y.P.D.; Methodology, Q.L., J.M.P. and Y.P.D.; Investigation, Q.L. and Y.P.D.; Writing – Original Draft, Q.L.; Writing – Review & Editing, J.M.P. and Y.P.D.; Funding Acquisition, Y.P.D.; Resources, J.M.P. and Y.P.D.; Supervision, Y.P.D.

## Reference

1. Shah VS, Meyerholz DK, Tang XX, Reznikov L, Abou Alaiwa M, Ernst SE, Karp PH, Wohlford-Lenane CL, Heilmann KP, Leidinger MR, Allen PD, Zabner J, McCray PB, Ostedgaard LS, Stoltz DA, Randak CO, Welsh MJ. 2016. Airway acidification initiates host defense abnormalities in cystic fibrosis mice. Science 351:503–507.

2. Pezzulo AA, Tang XX, Hoegger MJ, Abou Alaiwa MH, Ramachandran S, Moninger TO, Karp PH, Wohlford-Lenane CL, Haagsman HP, van Eijk M, Banfi B, Horswill AR, Stoltz DA, McCray PB, Jr., Welsh MJ, Zabner J. 2012. Reduced airway surface pH impairs bacterial killing in the porcine cystic fibrosis lung. Nature 487:109–13.

3. Zabner J, Smith JJ, Karp PH, Widdicombe JH, Welsh MJ. 1998. Loss of CFTR chloride channels alters salt absorption by cystic fibrosis airway epithelia in vitro. Mol Cell 2:397–403.

4. Silva Filho LV, Ferreira Fde A, Reis FJ, Britto MC, Levy CE, Clark O, Ribeiro JD. 2013. Pseudomonas aeruginosa infection in patients with cystic fibrosis: scientific evidence regarding clinical impact, diagnosis, and treatment. J Bras Pneumol 39:495–512.

5. Foundation CF. 2019. 2018 Annual Data Report Bethesda, Maryland. Cystic Fibrosis Foundation Patient Registry.

6. Lyczak JB, Cannon CL, Pier GB. 2002. Lung infections associated with cystic fibrosis. Clin Microbiol Rev 15:194–222.

7. Matsukawa M, Greenberg EP. 2004. Putative exopolysaccharide synthesis genes influence Pseudomonas aeruginosa biofilm development. J Bacteriol 186:4449–56.

8. Okshevsky M, Meyer RL. 2015. The role of extracellular DNA in the establishment, maintenance and perpetuation of bacterial biofilms. Crit Rev Microbiol 41:341–52.

9. Wilton M, Charron-Mazenod L, Moore R, Lewenza S. 2016. Extracellular DNA Acidifies Biofilms and Induces Aminoglycoside Resistance in Pseudomonas aeruginosa. Antimicrob Agents Chemother 60:544–53.

10. Lethem MI, James SL, Marriott C, Burke JF. 1990. The origin of DNA associated with mucus glycoproteins in cystic fibrosis sputum. Eur Respir J 3:19–23.

11. Chen JH, Stoltz DA, Karp PH, Ernst SE, Pezzulo AA, Moninger TO, Rector MV, Reznikov LR, Launspach JL, Chaloner K, Zabner J, Welsh MJ. 2010. Loss of anion transport without increased sodium absorption characterizes newborn porcine cystic fibrosis airway epithelia. Cell 143:911–23.

12. Mayer-Hamblett N, Kronmal RA, Gibson RL, Rosenfeld M, Retsch-Bogart G, Treggiari MM, Burns JL, Khan U, Ramsey BW, Investigators E. 2012. Initial Pseudomonas aeruginosa treatment failure is associated with exacerbations in cystic fibrosis. Pediatr Pulmonol 47:125–34.

13. Abou Alaiwa MH, Reznikov LR, Gansemer ND, Sheets KA, Horswill AR, Stoltz DA, Zabner J, Welsh MJ. 2014. pH modulates the activity and synergism of the airway surface liquid antimicrobials beta-defensin-3 and LL-37. Proc Natl Acad Sci U S A 111:18703–8.

14. Johansson J, Gudmundsson GH, Rottenberg ME, Berndt KD, Agerberth B. 1998. Conformation-dependent antibacterial activity of the naturally occurring human peptide LL-37. J Biol Chem 273:3718–24.

15. Nakayama K, Jia YX, Hirai H, Shinkawa M, Yamaya M, Sekizawa K, Sasaki H. 2002. Acid stimulation reduces bactericidal activity of surface liquid in cultured human airway epithelial cells. Am J Respir Cell Mol Biol 26:105–13.

16. Tang XX, Ostedgaard LS, Hoegger MJ, Moninger TO, Karp PH, McMenimen JD, Choudhury B, Varki A, Stoltz DA, Welsh MJ. 2016. Acidic pH increases airway surface liquid viscosity in cystic fibrosis. J Clin Invest 126:879–91.

17. Adewoye LO, Worobec EA. 1999. Multiple environmental factors regulate the expression of the carbohydrate-selective OprB porin of Pseudomonas aeruginosa. Can J Microbiol 45:1033–42.

18. Walton WG, Ahmad S, Little MS, Kim CS, Tyrrell J, Lin Q, Di YP, Tarran R, Redinbo MR. 2016. Structural Features Essential to the Antimicrobial Functions of Human SPLUNC1. Biochemistry 55:2979–91.

19. Di YP, Tkach AV, Yanamala N, Stanley S, Gao S, Shurin MR, Kisin ER, Kagan VE, Shvedova A. 2013. Dual Acute Pro-Inflammatory and Anti-Fibrotic Pulmonary Effects of SPLUNC1 After Exposure to Carbon Nanotubes. Am J Respir Cell Mol Biol doi:10.1165/rcmb.2012-0435OC.

20. Dobay O, Laub K, Stercz B, Keri A, Balazs B, Tothpal A, Kardos S, Jaikumpun P, Ruksakiet K, Quinton PM, Zsembery A. 2018. Bicarbonate Inhibits Bacterial Growth and Biofilm Formation of Prevalent Cystic Fibrosis Pathogens. Front Microbiol 9:2245.

21. Abou Alaiwa MH, Launspach JL, Grogan B, Carter S, Zabner J, Stoltz DA, Singh PK, McKone EF, Welsh MJ. 2018. Ivacaftor-induced sweat chloride reductions correlate with increases in airway surface liquid pH in cystic fibrosis. JCI Insight 3.

22. Whiteley M, Bangera MG, Bumgarner RE, Parsek MR, Teitzel GM, Lory S, Greenberg EP. 2001. Gene expression in Pseudomonas aeruginosa biofilms. Nature 413:860–4.

23. Bryan LE, Kwan S. 1983. Roles of ribosomal binding, membrane potential, and electron transport in bacterial uptake of streptomycin and gentamicin. Antimicrob Agents Chemother 23:835–45.

24. Bryan LE, Nicas T, Holloway BW, Crowther C. 1980. Aminoglycoside-resistant mutation of Pseudomonas aeruginosa defective in cytochrome c552 and nitrate reductase. Antimicrob Agents Chemother 17:71–9.

25. Bryan LE, Haraphongse R, Van den Elzen HM. 1976. Gentamicin resistance in clinical-isolates of Pseudomonas aeruginosa associated with diminished gentamicin accumulation and no detectable enzymatic modification. J Antibiot (Tokyo) 29:743–53.

26. MacLeod DL, Nelson LE, Shawar RM, Lin BB, Lockwood LG, Dirk JE, Miller GH, Burns JL, Garber RL. 2000. Aminoglycoside-resistance mechanisms for cystic fibrosis Pseudomonas aeruginosa isolates are unchanged by long-term, intermittent, inhaled tobramycin treatment. J Infect Dis 181:1180–4.

27. Rivera M, Hancock RE, Sawyer JG, Haug A, McGroarty EJ. 1988. Enhanced binding of polycationic antibiotics to lipopolysaccharide from an aminoglycoside-supersusceptible, tolA mutant strain of Pseudomonas aeruginosa. Antimicrob Agents Chemother 32:649–55.

28. Breedveld MW, Miller KJ. 1994. Cyclic beta-glucans of members of the family Rhizobiaceae. Microbiol Rev 58:145–61.

29. Mah TF, Pitts B, Pellock B, Walker GC, Stewart PS, O’Toole GA. 2003. A genetic basis for Pseudomonas aeruginosa biofilm antibiotic resistance. Nature 426:306–10.

30. Bhagwat AA, Gross KC, Tully RE, Keister DL. 1996. Beta-glucan synthesis in Bradyrhizobium japonicum: characterization of a new locus (ndvC) influencing beta-(1-->6) linkages. J Bacteriol 178:4635–42.

31. Deziel E, Lepine F, Milot S, Villemur R. 2003. rhlA is required for the production of a novel biosurfactant promoting swarming motility in Pseudomonas aeruginosa: 3-(3-hydroxyalkanoyloxy)alkanoic acids (HAAs), the precursors of rhamnolipids. Microbiology 149:2005–13.

32. McClure CD, Schiller NL. 1996. Inhibition of macrophage phagocytosis by Pseudomonas aeruginosa rhamnolipids in vitro and in vivo. Curr Microbiol 33:109–17.

33. Read RC, Roberts P, Munro N, Rutman A, Hastie A, Shryock T, Hall R, McDonald-Gibson W, Lund V, Taylor G, et al. 1992. Effect of Pseudomonas aeruginosa rhamnolipids on mucociliary transport and ciliary beating. J Appl Physiol (1985) 72:2271–7.

34. Huang B, Whitchurch CB, Mattick JS. 2003. FimX, a multidomain protein connecting environmental signals to twitching motility in Pseudomonas aeruginosa. J Bacteriol 185:7068–76.

35. Chiang P, Burrows LL. 2003. Biofilm formation by hyperpiliated mutants of Pseudomonas aeruginosa. J Bacteriol 185:2374–8.

36. O’Toole GA, Kolter R. 1998. Flagellar and twitching motility are necessary for Pseudomonas aeruginosa biofilm development. Mol Microbiol 30:295–304.

37. Barbieri JT, Sun J. 2004. Pseudomonas aeruginosa ExoS and ExoT. Rev Physiol Biochem Pharmacol 152:79–92.

38. Hauser AR. 2009. The type III secretion system of Pseudomonas aeruginosa: infection by injection. Nat Rev Microbiol 7:654–65.

39. Martin DW, Schurr MJ, Mudd MH, Deretic V. 1993. Differentiation of Pseudomonas aeruginosa into the alginate-producing form: inactivation of mucB causes conversion to mucoidy. Molecular Microbiology 9:497–506.

40. Boucher JC, Schurr MJ, Yu H, Rowen DW, Deretic V. 1997. Pseudomonas aeruginosa in cystic fibrosis: role of mucC in the regulation of alginate production and stress sensitivity. Microbiology 143 (Pt 11):3473–80.

41. Wozniak DJ, Wyckoff TJO, Starkey M, Keyser R, Azadi P, O’Toole GA, Parsek MR. 2003. Alginate is not a significant component of the extracellular polysaccharide matrix of PA14 and PAO1 Pseudomonas aeruginosa biofilms. Proc Natl Acad Sci U S A 100:7907–12.

42. Johnson L, Mulcahy H, Kanevets U, Shi Y, Lewenza S. 2012. Surface-localized spermidine protects the Pseudomonas aeruginosa outer membrane from antibiotic treatment and oxidative stress. J Bacteriol 194:813–26.

43. Ernst RK, Moskowitz SM, Emerson JC, Kraig GM, Adams KN, Harvey MD, Ramsey B, Speert DP, Burns JL, Miller SI. 2007. Unique lipid a modifications in Pseudomonas aeruginosa isolated from the airways of patients with cystic fibrosis. J Infect Dis 196:1088–92.

44. Schultz A, Puvvadi R, Borisov SM, Shaw NC, Klimant I, Berry LJ, Montgomery ST, Nguyen T, Kreda SM, Kicic A, Noble PB, Button B, Stick SM. 2017. Airway surface liquid pH is not acidic in children with cystic fibrosis. Nature Communications 8:1409.

45. McShane D, Davies JC, Davies MG, Bush A, Geddes DM, Alton EW. 2003. Airway surface pH in subjects with cystic fibrosis. Eur Respir J 21:37–42.

46. Hunter RC, Beveridge TJ. 2005. Application of a pH-sensitive fluoroprobe (C-SNARF-4) for pH microenvironment analysis in Pseudomonas aeruginosa biofilms. Appl Environ Microbiol 71:2501–10.

47. de los Rios A, Wierzchos J, Sancho LG, Ascaso C. 2003. Acid microenvironments in microbial biofilms of antarctic endolithic microecosystems. Environ Microbiol 5:231–7.

48. Hidalgo G, Burns A, Herz E, Hay AG, Houston PL, Wiesner U, Lion LW. 2009. Functional tomographic fluorescence imaging of pH microenvironments in microbial biofilms by use of silica nanoparticle sensors. Appl Environ Microbiol 75:7426–35.

49. Tate S, MacGregor G, Davis M, Innes JA, Greening AP. 2002. Airways in cystic fibrosis are acidified: detection by exhaled breath condensate. Thorax 57:926–9.

50. Ojoo JC, Mulrennan SA, Kastelik JA, Morice AH, Redington AE. 2005. Exhaled breath condensate pH and exhaled nitric oxide in allergic asthma and in cystic fibrosis. Thorax 60:22–6.

51. Smith EE, Buckley DG, Wu Z, Saenphimmachak C, Hoffman LR, D’Argenio DA, Miller SI, Ramsey BW, Speert DP, Moskowitz SM, Burns JL, Kaul R, Olson MV. 2006. Genetic adaptation by Pseudomonas aeruginosa to the airways of cystic fibrosis patients. Proc Natl Acad Sci U S A 103:8487–92.

52. Bragonzi A, Paroni M, Nonis A, Cramer N, Montanari S, Rejman J, Di Serio C, Doring G, Tummler B. 2009. Pseudomonas aeruginosa microevolution during cystic fibrosis lung infection establishes clones with adapted virulence. Am J Respir Crit Care Med 180:138–45.

53. Rahme LG, Stevens EJ, Wolfort SF, Shao J, Tompkins RG, Ausubel FM. 1995. Common virulence factors for bacterial pathogenicity in plants and animals. Science 268:1899–902.

54. O’Toole GA. 2011. Microtiter dish biofilm formation assay. J Vis Exp doi:10.3791/2437.

55. Poltak SR, Cooper VS. 2011. Ecological succession in long-term experimentally evolved biofilms produces synergistic communities. ISME J 5:369–78.

56. Cooper VS. 2018. Experimental Evolution as a High-Throughput Screen for Genetic Adaptations. mSphere 3.

57. Jiang S, Deslouches B, Chen C, Di ME, Di YP. 2019. Antibacterial Properties and Efficacy of a Novel SPLUNC1-Derived Antimicrobial Peptide, alpha4-Short, in a Murine Model of Respiratory Infection. MBio 10.

58. Liu Y, Bartlett JA, Di ME, Bomberger JM, Chan YR, Gakhar L, Mallampalli RK, McCray PB, Jr., Di YP. 2013. SPLUNC1/BPIFA1 contributes to pulmonary host defense against Klebsiella pneumoniae respiratory infection. Am J Pathol 182:1519–31.

59. Liu Y, Di ME, Chu HW, Liu X, Wang L, Wenzel S, Di YP. 2013. Increased susceptibility to pulmonary Pseudomonas infection in Splunc1 knockout mice. J Immunol 191:4259–68.

60. Cury JA, Koo H. 2007. Extraction and purification of total RNA from Streptococcus mutans biofilms. Anal Biochem 365:208–14.

61. Lin Q, Deslouches B, Montelaro RC, Di YP. 2018. Prevention of ESKAPE pathogen biofilm formation by antimicrobial peptides WLBU2 and LL37. Int J Antimicrob Agents 52:667–672.

62. Casciaro B, Lin Q, Afonin S, Loffredo MR, de Turris V, Middel V, Ulrich AS, Di YP, Mangoni ML. 2019. Inhibition of Pseudomonas aeruginosa biofilm formation and expression of virulence genes by selective epimerization in the peptide Esculentin-1a(1-21)NH2. FEBS J 286:3874–3891.

63. Lin Q, Di YP. 2020. Determination and Quantification of Bacterial Virulent Gene Expression Using Quantitative Real-Time PCR. Methods Mol Biol 2102:177–193.

