## Supplemental data for "Cystic Fibrosis Acidic Microenvironment Determines Antibiotic Susceptibility and Biofilm Formation of *Pseudomonas aeruginosa*"

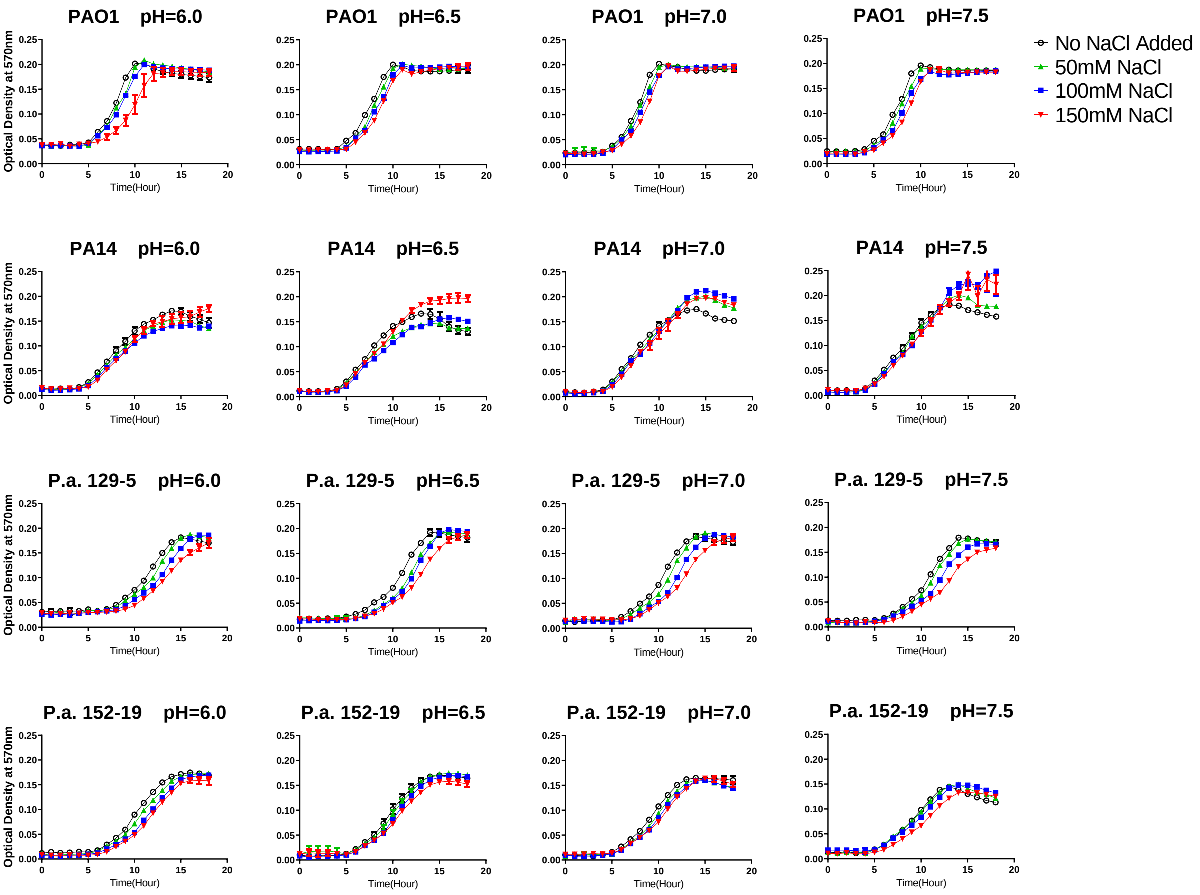


**Fig. S1. Adjusted salt and pH conditions do not significantly affect the proliferation rate of planktonic *P. aeruginosa*.** PAO1, PA14, *P.a.*129-5 and *P.a.*152-19 were inoculated in 10% TSB (n=3). The salt concentration of the culture media was adjusted by addition of 50, 100 and 150mM NaCl. No NaCl group (pH=7.5) was used as control. Optical density at 570nm was measured every hour for 18 hours at 37°C in a microplate reader.

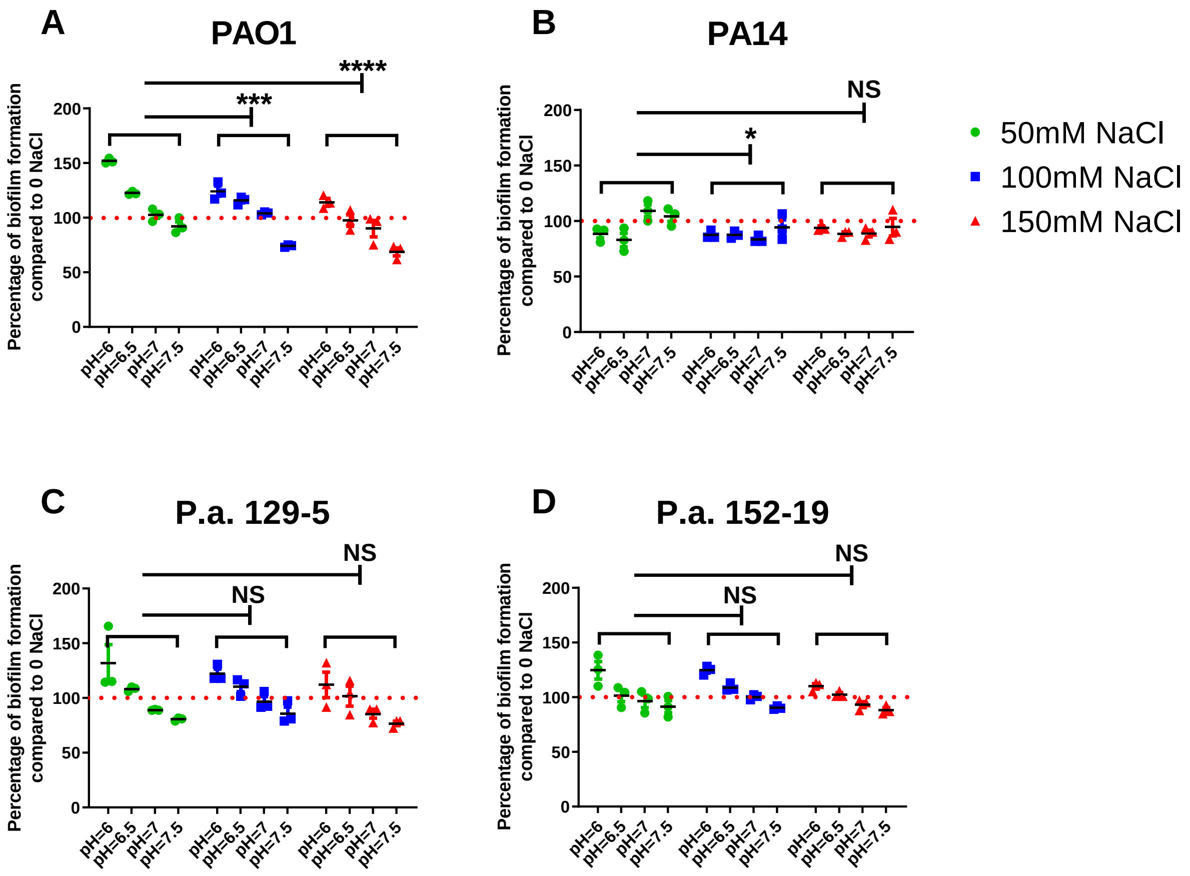


**Fig. S2. Elevated NaCl concentrations have minimal effect on *P. aeruginosa* biofilm formation but acidic pH promotes *P. aeruginosa* biofilm formation.** (**A** to **D**) Effect of NaCl (50, 100 and 150mM) and pH (6.0, 6.5, 7.0 and 7.5) on *P. aeruginosa* biofilm formation compared to positive controls (n=3). Bacteria were incubated for 3 hours at 37°C in 96-well microplates. The crystal violet staining method was used to quantify the biofilm/biomass attachment. The red dotted lines denote the (+) control of biofilm formation at pH 7.5 without additional NaCl. Data are mean ± SEM. Two-way ANOVA was used for statistical analysis. ***p*<0.01; ****p*<0.001; *****p* <0.0001; NS: not significant.

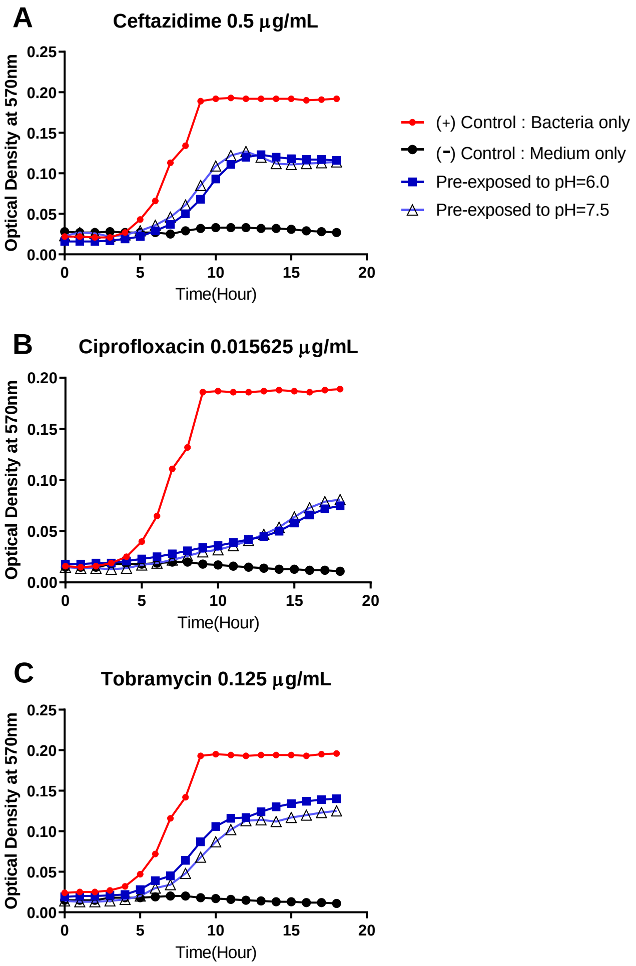


**Fig. S3. Antibiotics restore their antimicrobial activity in physiological pH condition after prior exposure to acidic environment.** Antibiotics (ceftazidime, ciprofloxacin and tobramycin) were pre-exposed to pH = 6.0 or 7.5 in 10% TSB for 5 hours, the pH for all exposed antibiotics was then re-adjusted back to 7.5 before the antibiotics were added into planktonic *P. aeruginosa* (PAO1) culture for GIA at pH 7.5. The applied treatment concentration for each antibiotic was selected using less than their respective MIC to demonstrate the partial inhibition of bacterial growth. Optical density at 570nm was measured every hour for 18 hours at 37°C in a microplate reader. Results are mean ± SD from two independent experiments with bacteria grew in duplicates for each condition.

### Ouabain and DMSO vehicle are not bactericidal to PAO1

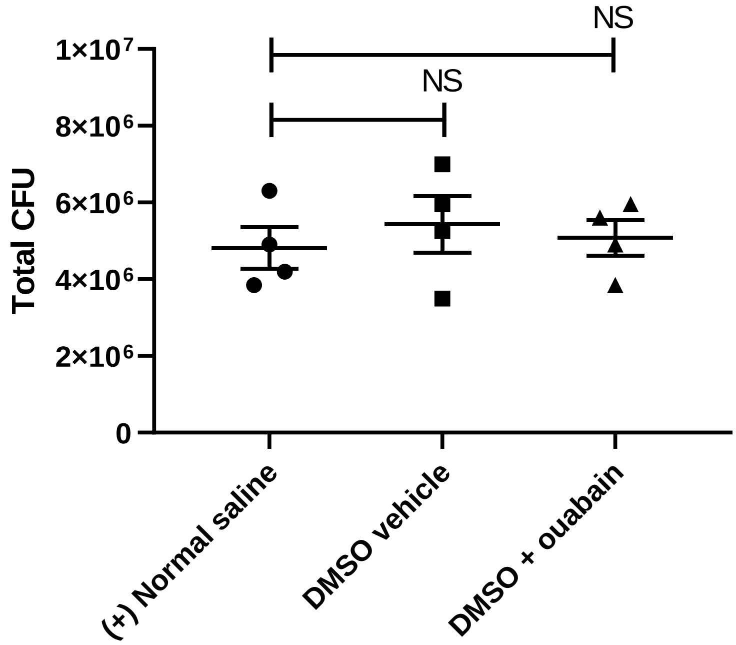


**Fig. S4. Ouabain treatment was not bactericidal to PAO1.** A total volume of 50µL of 2 x $10^8$ CFU/mL PAO1 and 20µL of 20µM Ouabain or DMSO (solvent control) were incubated in normal saline for 5 hours at 37°C (n=4). Data are mean ± SEM from two repeated experiments. Student's *t* tests were used for statistical analysis. NS: not significant.

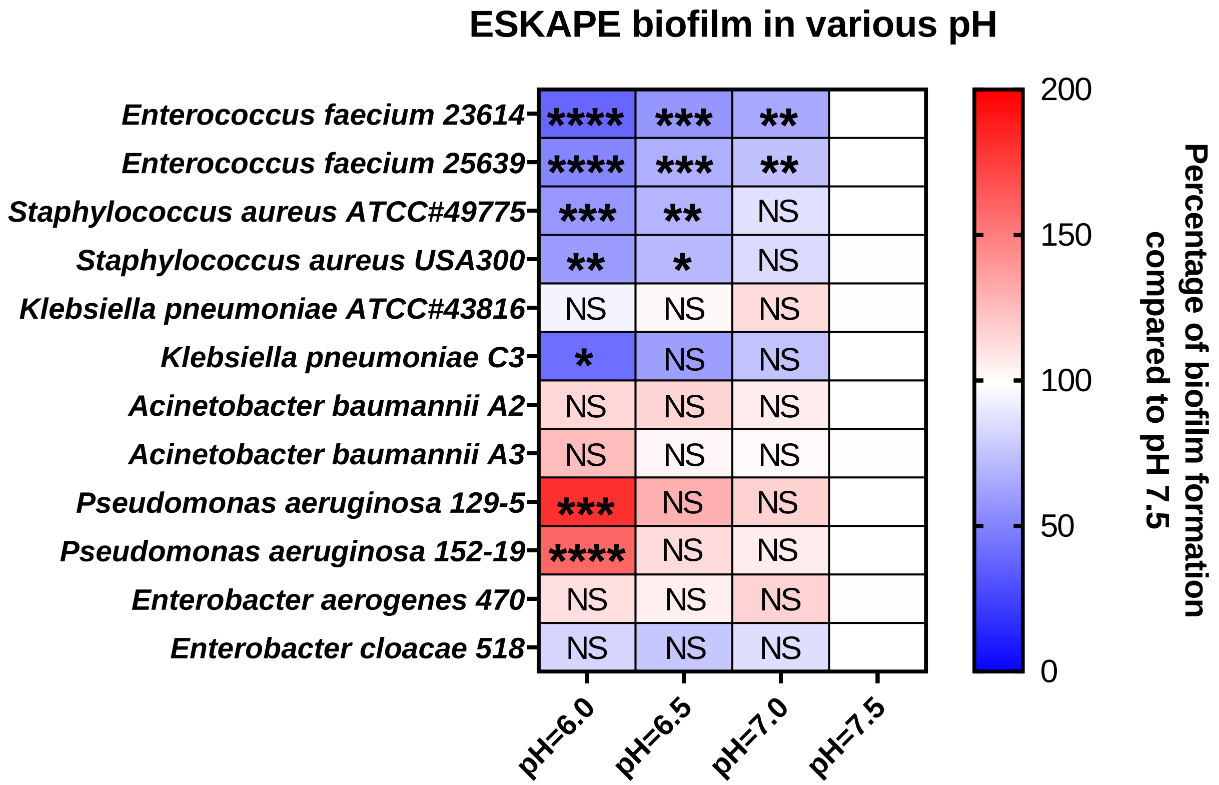


**Fig. S5. Effects of acidic pH on ESKAPE pathogen biofilm formation**. ESKAPE pathogen biofilm was quantified by the crystal violet method after 3 hours of incubation. The pH 7.5 biofilm formation group was served as a control. The color scale bars represent percentage of biofilm formation compared to the respective control groups. Red scale: increased biofilm formation (100% - 200%); white: no change (100%); blue scale: decreased biofilm formation (0 – 100%). Data are mean ± SEM (n=3). One-way ANOVA test was performed to compare pH 6.0, 6.5, and 7.0 biofilm formation to pH 7.5. **p*<0.05; ***p* <0.01; ****p* <0.001; *****p* <0.0001; NS: not significant.

**Supplementary Table 1. Effects of acidic pH on *P. aeruginosa* antibiotic resistance in MIC changes.** MIC data were acquired using the GIA assay. Results are averaged MICs from two independent GIA experiments.

| Ceftazidime MIC (µg/mL) | | | | |
|---|---|---|---|---|
| NO. | Strain | pH 6.5 average MIC | pH 7.5 average MIC | MIC fold increase from pH 6.5 to pH |
| 1 | PAO1 | 1.875 | 0.9375 | 2 |
| 2 | PA14 | 2.5 | 1.25 | 2 |
| 3 | 116-37 | 2.5 | 0.9375 | 3 |
| 4 | 71-75 | 0.625 | 0.15625 | 4 |
| 5 | KK1 | 2.5 | 1.25 | 2 |
| 6 | AA43 | 5 | 2.5 | 2 |
| 7 | 40-21 | 2.5 | 1.25 | 2 |
| 8 | 151-40 | 5 | 2.5 | 2 |
| 9 | 129-5 | 5 | 1.25 | 4 |
| 10 | 109-10 | 2.5 | 1.25 | 2 |
| 11 | 480-1 | 0.625 | 0.15625 | 4 |
| 12 | 82-9 | 2.5 | 1.25 | 2 |
| 13 | 152-19 | 5 | 2.5 | 2 |
| 14 | 80-37 | 2.5 | 1.25 | 2 |

| Ciprofloxacin MIC (µg/mL) | | | | |
|---|---|---|---|---|
| NO. | Strain | pH 6.5 average MIC | pH 7.5 average MIC | MIC fold increase from pH 6.5 to pH |
| 1 | PAO1 | 0.039063 | 0.039063 | 1 |
| 2 | PA14 | 0.07813 | 0.039063 | 2 |
| 3 | 116-37 | 0.07813 | 0.039063 | 2 |
| 4 | 71-75 | 0.15625 | 0.117188 | 1 |
| 5 | KK1 | 0.03906 | 0.039063 | 1 |
| 6 | AA43 | 0.15625 | 0.078125 | 2 |
| 7 | 40-21 | 0.07813 | 0.039063 | 2 |
| 8 | 151-40 | 0.9375 | 0.625 | 2 |
| 9 | 129-5 | 0.07813 | 0.039063 | 2 |
| 10 | 109-10 | 0.15625 | 0.058594 | 3 |
| 11 | 480-1 | 0.014648 | 0.014649 | 1 |
| 12 | 82-9 | 0.07813 | 0.039063 | 2 |
| 13 | 152-19 | 0.07813 | 0.039063 | 2 |
| 14 | 80-37 | 0.039063 | 0.039063 | 1 |

| Tobramycin MIC (µg/mL) | | | | |
|---|---|---|---|---|
| NO. | Strain | pH 6.5 average MIC | pH 7.5 average MIC | MIC fold increase from pH 6.5 to pH |
| 1 | PAO1 | 2.5 | 0.9375 | 3 |
| 2 | PA14 | 3.75 | 1.25 | 3 |
| 3 | 116-37 | 1.25 | 0.3125 | 4 |
| 4 | 71-75 | 2.5 | 0.625 | 4 |
| 5 | KK1 | 5 | 1.25 | 4 |
| 6 | AA43 | 10 | 1.875 | 5 |
| 7 | 40-21 | 1.25 | 0.3125 | 4 |
| 8 | 151-40 | 2.5 | 1.25 | 2 |
| 9 | 129-5 | 0.625 | 0.625 | 1 |
| 10 | 109-10 | 10 | 2.5 | 4 |
| 11 | 480-1 | 0.9375 | 0.3125 | 3 |
| 12 | 82-9 | 5 | 0.625 | 8 |
| 13 | 152-19 | 1.25 | 0.625 | 2 |
| 14 | 80-37 | 5 | 0.9375 | 5 |

**Supplementary Table 2. List of primers used for real time qPCR.**

| Gene | Primer sequence | |
|---|---|---|
| *tolA* | Forward | 5’ - GCG TAA TGG AAT GAG CGT AGA A -3’ |
| | Reverse | 5’ - CGA ACT GTC GAA AGG CTT GT - 3’ |
| *ndvB* | Forward | 5’ - TGT GGA TCG CCT ACG ACT A - 3’ |
| | Reverse | 5’ - CGG TGA ACA GCA CGA TGA - 3’ |
| *mucB* | Forward | 5’-TTT GCT TGG CAG CCT GAT-3’ |
| | Reverse | 5’-GGT GCC TTG GAA ACT GTT CT-3’ |
| *exoS* | Forward | 5’-GAC AGG CTG AAC AGG TAG TG-3’ |
| | Reverse | 5’-TTC AGG GAG GTG GAG AGA TAG-3’ |
| *exoT* | Forward | 5’-GCT GAA CAG GTC GTG AAG A-3’ |
| | Reverse | 5’-CCG GGA GGT GGA GAG ATA G-3’ |
| *fimT* | Forward | 5’-CGC TTG CAA AGA AGG AAA GG-3’ |
| | Reverse | 5’-CCT TCC GCA GAG CAG AAA-3’ |
| *fimX* | Forward | 5’-CCT GGC CTA TAT CCA TCT CAA C-3’ |
| | Reverse | 5’-ACT GTT CAC GCA TCA GTC C-3’ |
| *rhlA* | Forward | 5’-CGA GAC CGT CGG CAA ATA C-3’ |
| | Reverse | 5’-GCA CCT GGT CGA TGT GAA A-3’ |
| *rhlB* | Forward | 5’-CTC ACG AGA AGT ACG GGA TTC-3’ |
| | Reverse | 5’-CTC GGG CAC GTT GAA CT-3’ |
| *rplU* * | Forward | 5’-CGC AGT GAT TGT TAC CGG TG-3’ |
| | Reverse | 5’-AGG CCT GAA TGC CGG TGA TC-3’ |

*The constitutively expressed *rplU* gene served as a reference gene.
